# A naturalistic assay for measuring behavioral responses to aversive stimuli at millisecond timescale

**DOI:** 10.1101/161885

**Authors:** Carl E. Schoonover, Andrew J. P. Fink, Richard Axel

## Abstract

We have designed a Virtual Burrow Assay (VBA) to detect the behavioral responses of head-fixed mice to aversive stimuli. We demonstrate its suitability for measuring novelty detection as well as aversion to both conditioned and innately aversive cues. The VBA simulates a scenario in which a mouse, poised at the threshold of its burrow, evaluates whether to remain exposed to potential threats outside or to retreat inside an enclosure. When presented with aversive stimuli, mice exhibit a stereotyped retreat whose onset is determined by measuring the position of a moveable burrow. This withdrawal, which requires no training, is characterized by an abrupt transition that unfolds within milliseconds—a timescale similar to that of neuronal dynamics, permitting direct comparison between the two. The assay is compatible with standard electrophysiological and optical methods for measuring and perturbing neuronal activity.

## Introduction

Traditional assays of aversion measure behavioral responses that extend over many seconds, such as time spent freezing during a specified epoch (Bouton and Bolles, 1980) or conditioned suppression (Estes and Skinner, 1941). The flight from danger, in contrast, is a stereotyped behavioral motif whose onset consists of a rapid transition in behavioral state (Blanchard et al., 1998; De Franceschi et al., 2016; Domenici and Blake, 1991; Walther, 1969; Yilmaz and Meister, 2013). Mice, for example, initiate flight within 250 msec of the presentation of a looming visual stimulus (De Franceschi et al., 2016; Yilmaz and Meister, 2013). The capacity to precisely measure the onset of behavioral responses to aversive cues would permit comparison of transitions in behavioral state and transitions in neuronal activity. Here we describe a Virtual Burrow Assay for head-fixed mice that detects the temporally precise onset of responses to aversive stimuli by exploiting a rapid and stereotyped motor sequence.

The Virtual Burrow Assay (VBA) recapitulates in the laboratory the behavioral contingencies of a mouse poised at the threshold of its burrow. In the wild, mice in this state must evaluate whether it is safe to exit, or whether a perceived threat warrants hasty retreat to the safety of the burrow (Birke et al., 1985; Blanchard and Blanchard, 2008; Blanchard and Blanchard, 1989). The VBA captures behavioral responses with millisecond precision, exploits a stereotyped behavior that requires no training, and is compatible with standard electrophysiological and optical methods for measuring and perturbing neuronal activity. Retreat in the VBA marks an abrupt behavioral transition that unfolds at a timescale similar to that of neuronal dynamics. This precise timestamp permits alignment of behavioral state with underlying neuronal activity, as does the eye saccade in primate neurophysiology (Roitman and Shadlen, 2002).

### Rapid and reliable ingress in response to air puff

The VBA consists of a tube enclosure (virtual burrow), constrained to slide back and forth along the anterior-posterior axis of the body of a head-fixed mouse (Figure 1). When placed inside the virtual burrow, mice invariably attempt to enter the tube as far as possible, pulling it to an “ingress” position, where they remain during an initial acclimation period. In preparation for each trial, a tether pulls the tube away from the body to an “egress” position, forcing the animal to exit the burrow. After initially resisting the retraction of the burrow, mice eventually maintain this exposed, egress position voluntarily. The tether is then slackened to permit free ingress and stimuli are presented while the mouse is in full control of the position of the virtual burrow (Figure 1D). The assay measures the position of the burrow on a millisecond timescale and detects the precise timing of the transition from egress (Figure 1A, second from right) to ingress (Figure 1A, far right).

**Figure 1.**
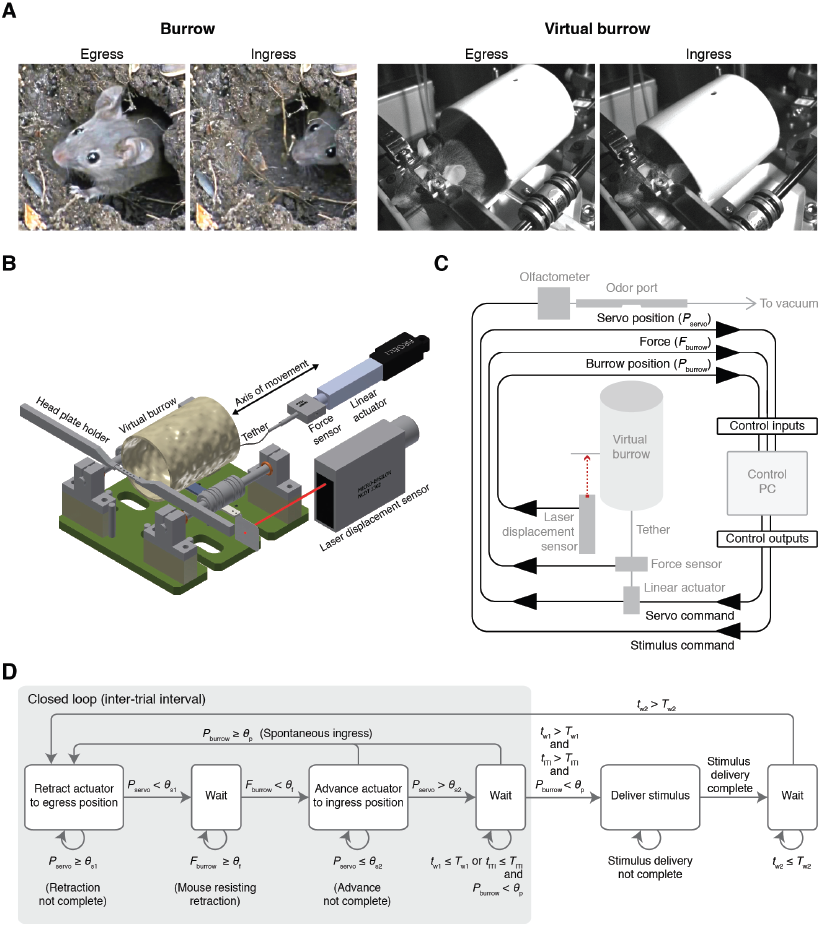
The Virtual Burrow Assay. A.Left, mouse in the wild exiting (egress) and entering (ingress) its burrow (stills courtesy of Misterduncan,YouTube, March 25, 2009). Right, head-fixed mouse exiting (egress) and entering (ingress) the virtual burrow.B.Instrument diagram. A head-fixed mouse is secured by a headplate holder and stands inside a virtual burrow (cardboard or 3D-printed tube). Alinear actuator can retract the burrow along a single axis of movement, constrained by a pair of near-frictionless air bearings. The mass of the tube and bearing system is comparable to the weight of an adult mouse (~30 g). A laser displacement sensor measures burrow position and a force sensor measuresA.the force generated by the animal when pulling via the tether against the linear actuator (Rendering courtesy of Tanya Tabachnik, Advanced Instrumentation, Zuckerman Mind Brain Behavior Institute). C. Schematic of the virtual burrow control system. During the intertrial interval (ITI), the linear actuator retracts the burrow to the egress position. Once the animal’s initial resistance subsides, as measured by the force sensor, the linear actuator advances, slackening the tether and freeing the animal to ingress. Trial initiation occurs once the animal has freely maintained the egress position for at least 10 sec. In the event of premature ingress, the trial is aborted and the linear actuator retracts. Control inputs and outputs, black. Devices, grey. D. Finite state machine diagram. During the ITI the burrow is retracted to the egress position by the linear actuator (“Retract actuator to egress position”). The retraction is complete once the position of the servo motor (*P*_servo_) is less than the specified retraction position (*θ*_s1_). As long as the force against the tether (*F*_burrow_) exceeds a preset threshold (*θ*_f_), the linear actuator remains in the retracted position (“Wait”). Once *F*_burrow_ < *θ*_f_, the linear actuator advances to the ingress position (“Advance actuator to ingress position”), slackening the tether. Once *P*_servo_ reaches the slacked position (*θ*_s2_), the system waits (“Wait”). If at any point the position of the burrow (*P*_burrow_) exceeds a specified threshold (*θ*_p_), the burrow is again retracted and the system returns to the initial state. If, however, the animal remains in the egress position and *P*_burrow_ < θ_S_ for a duration (*t*_ITI_) exceeding both the ITI (*T*_ITI_) and an enforced delay (*T*_w1_) following advance of the linear actuator to the ingress position (*t*_w1_), then the ITI concludes and a stimulus is delivered (“Deliver stimulus”). Throughout the period of stimulus delivery, the linear actuator remains in the advanced position with the animal in complete control of burrow position. Following stimulus delivery (*t*_w2_), once a second delay period (*T*_w2_) has elapsed, the system returns to the initial state and the linear actuator retracts the burrow.

We first determined whether an innately aversive stimulus, such as air puff, induces ingress, and if so, the degree to which this response is stereotyped and rapid (Figure 2A). Strong air puff delivered to the snout (80 PSI, 2-mm distance) elicited short latency, rapid ingress in all mice tested on all trials (n = 3 mice, 15 trials per mouse) (Figure 2B,C; Supplementary Video 1). Animals generated this behavior by pulling the burrow up to the ingress position in a coordinated, simultaneous movement of their fore- and hind-limbs. The latency to ingress varied little across 15 consecutive trials within each animal, but considerably across animals (Figure 2C). In contrast, a weak air puff (2 PSI, 15-cm distance) rarely elicited ingress and instead caused the animal to flinch. This apparent startle response (Davis, 1984) was visible as a transient change in burrow position clearly distinct from the ongoing movement of the burrow caused by the animal’s breathing (Figure 2D). The reliability and low latency of the response to strong air puff, together with the fact that no training is required, suggest that ingress in the VBA exploits an innate, highly stereotyped behavioral program. These results demonstrate that this assay is capable of capturing the fine temporal properties of this behavior.

**Figure 2.**
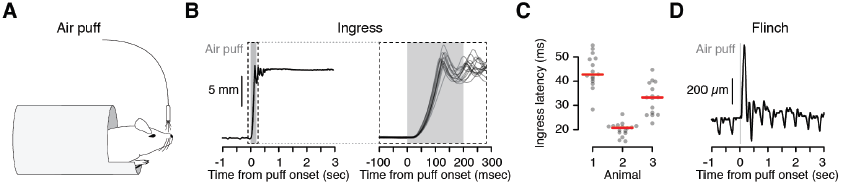
Reliable, short-latency ingress to noxious air puff. Diagram of experimental set up. With the mouse head-fixed within the virtual burrow, an air puff stimulus is delivered to the nose via a blunt syringe needle. B. Left, burrow position as a function of time showing a single ingress in response to a strong air puff (grey box, 200 msec, 80 psi). Upward deflections correspond to burrow movement towards the animal’s body (ingress). Downward deflections correspond to burrow movement away from the animal’s body (egress). Upward going, high-amplitude, sustained deflection corresponds to ingress following air puff. Right, 15 ingress responses from a single animal to 15 air puffs at high temporal resolution. Dashed box at left demarcates epoch in which scaling is expanded at right. C. Latency to ingress onset in three animals. Individual trials, grey points. Median, red line. D. Startle-like flinch in response to light air puff. Downward going, approximately 2-Hz oscillations correspond to the animal’s breathing cycle. Upward going low-amplitude, transient deflection corresponds to startle in response to air puff (grey box, 20 msec, 2 psi) directed towards animal’s nose.

### Ingress in response to a looming visual stimulus

We next evaluated how behavior in the VBA compares to the behavior of freely moving animals by presenting stimuli known to trigger specific behavioral responses (De Franceschi et al., 2016; Yilmaz and Meister, 2013). We employed a visual “looming” stimulus that elicits flight in freely moving mice (De Franceschi et al., 2016; Yilmaz and Meister, 2013) (Figure 3A) and observed that mice ingressed on 80% of trials in response to an expanding black disk displayed above their heads (Figure 3A, top left, B, “loom”, C, orange, and D, left; Supplementary Video 2). This response did not habituate, in contrast to the behavior of freely moving mice (De Franceschi et al., 2016). Presentation of either a contracting black disk (Figure 3A, top middle, B, “recede”, C, pink, and D, middle) or a small black disk sweeping across the visual field (Figure 3A, top right, B, “sweep”, C, green, and D, right) elicited ingress on only 0% or 20% of trials, respectively. Again, this observation is consistent with the behavior of freely moving mice, which do not exhibit flight in response to these stimuli (De Franceschi et al., 2016). These data show that a visual stimulus that selectively elicits flight in feely moving mice selectively elicits ingress in the VBA.

**Figure 3.**
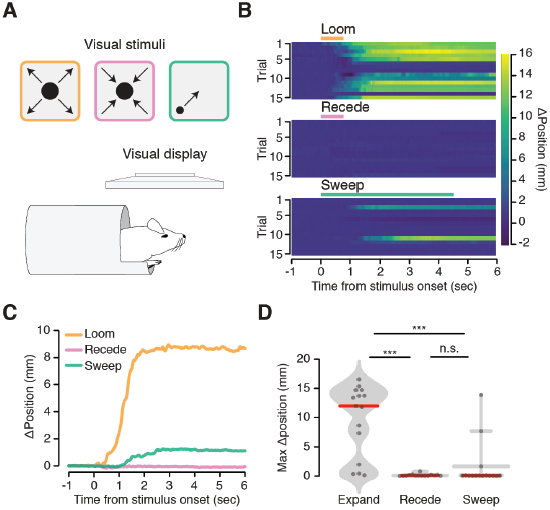
Flight-inducing visual stimuli selectively elicit ingress. An expanding black disk (left), a contracting black disk (middle), and a sweeping black disk of constant size (right), presented on a visual display positioned directly over a mouse head-fixed within the virtual burrow (bottom). B. Responses to three visual stimuli (n = 9 mice, 3 per condition, 5 trials each): Expanding (“loom”), disk widening from 2° to 50° over 250 msec, holding the 50° disk for 500 msec; Contracting (“recede”), disk diminishing from 50° to 2° over 250 msec, holding the 2° disk for 500 msec; Sweeping (“sweep”), 5° disk sweeping smoothly across the diagonal of the screen at a rate of 21°/sec (De Franceschi et al., 2016). Color map corresponds to change in burrow position (ΔPosition) with respect to baseline. C. Mean change in burrow position per condition across all animals and all trials. D. Maximum change in burrow position in the 6 sec following stimulus onset per condition across all animals and all trials. Ingress was defined as a maximum displacement of the burrow relative to the pre-stimulus baseline position > 0.85 mm during 5 sec following stimulus onset. The empirical likelihood of ingress was 0.80, 0.00 and 0.20 for loom, recede, and sweep, respectively (9 mice total, 3 mice per stimulus condition, 5 trials per mouse). Individual trials, grey points. Normalized, smoothed histogram, light grey shading. Median, red line. A two-proportion z-test was employed to evaluate whether the probability of ingress differs significantly across stimulus conditions. ***indicates p < 0.001, n.s. indicates p ≥ 0.05.

### Novel odor stimuli elicit ingress

We found that when presented with novel, neutral odorants mice initially ingress, but the response largely habituates after the first three trials (Odor 1, Figure 4A-C, first 15 blocks, and Figure 4D, left panel). We asked whether this behavior reflects nonspecific habituation to all olfactory stimuli or whether it is stimulus selective. After 15 presentations of Odor 1 we introduced a second, novel, neutral odor (Odor 2, 4B,C blocks 16-30, Odor 1 and Odor 2 presented in pseudorandom order within each block) and found that while mice selectively ingressed in response to Odor 2 during early trials, they remained unresponsive to the familiar odor. Similarly, after 15 blocks of Odors 1 and 2, we introduced a third, novel, neutral odor (Odor 3, Figure 4B,C, blocks 31-45, Odor 1, Odor 2, and Odor 3 presented in pseudorandom order within each block) and again observed ingress in response to the novel odor during early trials but not to the familiar ones. We found that the probability of ingress on the first three presentations of a given odor was significantly higher than the probability of ingress on later presentations of that same odor for all odors tested (Figure 4D). These data indicate that the VBA can be employed to detect selective responses to novel stimuli.

**Figure 4.**
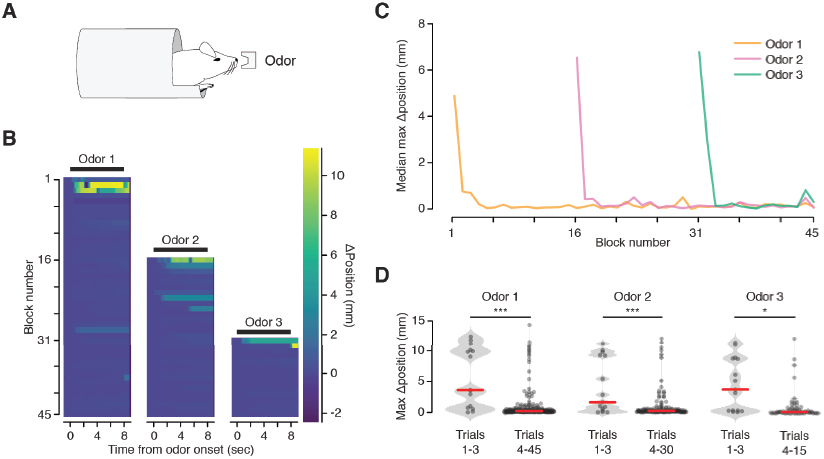
Habituation to novel stimuli. Three odor stimuli were delivered to mice within the virtual burrow assay. B. Habituating responses to sequential presentation of novel stimuli (Cooke et al. 2015) from a representative mouse. Color map corresponds to change in burrow position with respect to baseline. Black lines indicate odor stimulus epoch (8-sec duration).C. Median value across mice of maximum change in burrow position, per odor condition, per block. D. Maximum change in burrow position for each odor during the first three trials (left) and all later trials (right), pooled across animals. The probability of ingress (pingress) for all odors on the first three and all subsequent trials, Odor 1: p_ingress_ (trials 1-3) = 0.80, p_ingress_ (trials 4-45) = 0.18; Odor 2: p_ingress_ (trials 1-3) = 0.67, p_ingress_ (trials 4-30) = 0.20. Odor 3:p_ingress_ (trials 1-3) = 0.60, p_ingress_ (trials 4-15) = 0.23. Individual trials, grey points. Normalized, smoothed histogram, light grey shading. Median, red line. A two-proportion z-test on ingress probability pooled across all mice (n = 5) and all trials was employed to evaluate whether the probability of ingress differed significantly; ingress defined as maximum displacement > 0.75 mm during the 8-sec stimulus epoch. *indicates p < 0.05, ***indicates p < 0.001.

### Aversively conditioned odors selectively elicit ingress

We next determined whether the VBA can measure responses to aversively conditioned stimuli. We employed a classical conditioning paradigm in which on day 1 (Pre-test) we measured responses in the VBA to three neutral odorants. On day 2 (Conditioning) we placed the mice in a fear conditioning box and paired one odor with foot shock (CS+) and presented a second odor without foot shock (CS-); the third odor presented during pre-test (O3) was never presented on day 2. On day 3 (Test) we measured responses in the VBA to all three odor stimuli (Figure 5A).

**Figure 5.**
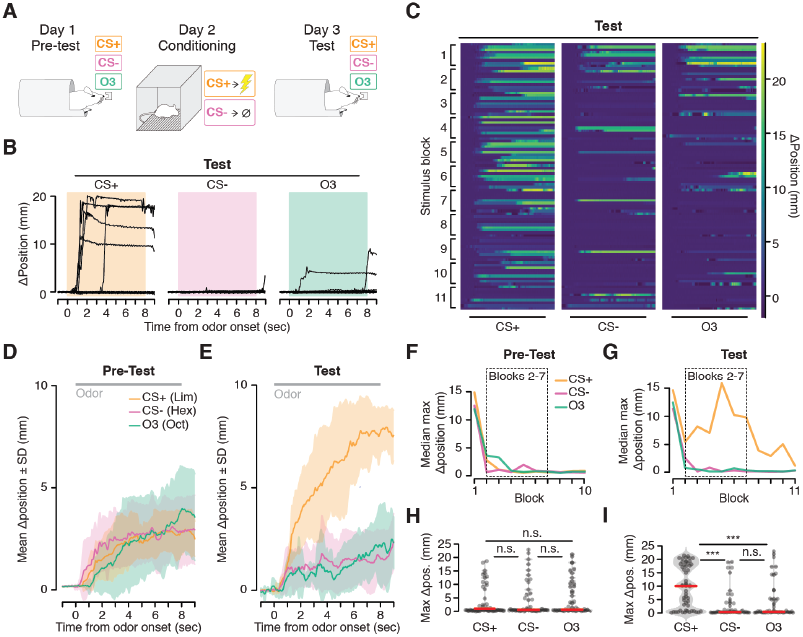
Ingress in response to aversively conditioned odor stimuli. A. Three odor stimuli were presented to mice head fixed in the VBA on day 1 (Pre-test) and day 3 (Test). On day 2 (Conditioning), animals were placed in a fear conditioning box and two of the odor stimuli were presented: a CS+ odor, paired with shock, and a CS- odor, never paired with shock. B. Change in burrow position relative to pre-stimulus baseline on individual trials after odor-shock conditioning from a representative mouse. Colored box demarcates odor stimulus epoch. CS+: paired with shock; CS-: unpaired with shock; Odor 3 (O3): not presented in fear conditioning box. C. Change in burrow position relative to pre-stimulus baseline (color map), ordered by mouse within each stimulus block. Lines at bottom indicates odor stimulus epoch (8-sec duration). D, E. Mean change in burrow position during Pre-test (D) and Test (E) relative to pre-stimulus baseline per odor condition during stimulus blocks 2 - 7 (shading indicates ±1 standard deviation, n = 9 mice). Grey line at top corresponds to odor stimulus epoch. F, G. Median value across mice of maximum change in burrow position, per odor condition, per block during Pre-test (F) and Test (G). Dashed boxes demarcate blocks 2 - 7, used to obtain mean responses and to perform statistical tests. H, I. Maximum change in burrow position during the odor stimulus, per condition across all animals and all trials during Pre-test (H) and Test (I). Individual trials, grey points. Normalized, smoothed histogram, light grey shading. Median, red line. The probability of ingress for each odor stimulus during Test: CS+: p_ingress_ = 0.76; CS-: p_ingress_ = 0.35; O3: p_ingress_ = 0.39; pingress(CS+) > p_ingress_ (CS-) p< 0.001; p_ingress_ (CS+) > p_ingress_ (O3) p < 0.001, two-proportion z-test on ingress probability pooled across all mice on blocks 2 - 7; ingress defined as maximum displacement > 0.75 mduring the 8-sec stimulus. ^***^ indicates p< 0.001, n.s. indicates p > 0.05.

We found that mice ingressed in response to all three odor stimuli during the first block of Pre-test (Figure 5F, block 1) and habituated over subsequent presentations (Figure 5D,F,H), consistent with our observation that animals ingress in response to novel odors (Figure 4). Following odor-shock pairing, we found that the likelihood of ingress was markedly higher in response to the CS+ than to the two stimuli not paired with shock (CS+ 76%, CS- 35%, O3 39%, Figure 5B,C,E,G,I). Moreover, mice ingressed markedly more to the CS+ during Test than during Pre-test (Test 76%, Pre-test 37%, Figure 5D versus E, F versus G, and H versus I). All nine mice tested exhibited a greater likelihood of ingress to CS+ than to CS- or O3; in eight out of nine this effect was significant (p < 0.05, two-proportion z-test on trials 2 - 7 to mitigate the effects of novelty observed on the first trial and extinction observed on the last three trials). This result is robust to the choice of ingress threshold over a range of 0.5 to 10 mm (Figure 5—figure supplement 1). Thus the VBA can detect stimulus-selective conditioned responses following aversive conditioning.

A subset of mice exhibited a second behavioral response specific to the CS+ following conditioning: an oscillation in burrow position (Figure 5—figure supplement 2). The high frequency of this oscillatory response distinguished it from the lower frequency oscillations that correspond to the animal’s breathing cycle (Figure 5—figure supplement 2, top and middle traces versus bottom trace; see also Figure 2D). Simultaneous video recording (not shown) indicated that it is instead associated with trembling of the animal’s body. This trembling sometimes preceded ingress by several seconds (Figure 5—figure supplement 2, top trace), or occurred on ingress-free trials(Figure 5—figure supplement 2, middle trace), and was selective for the CS+ stimulus following conditioning.

## Discussion

We have designed an assay to measure flight-like responses to both conditioned and innately aversive stimuli in head-fixed mice. The Virtual Burrow Assay captures a facet of the mouse *Umwelt* (the environment as it is experienced by members of a species) (von Uexküll, 1957) by simulating the scenario in which the animal is poised at the threshold of its burrow and evaluates whether to remain exposed to potential threats outside or to retreat inside the enclosure.

Looming stimuli that selectively evoke flight in freely moving mice (De Franceschi et al., 2016; Yilmaz and Meister, 2013) also selectively elicit ingress in head-fixed mice in the VBA (Figure 3). Noxious air puffs delivered to the nose invariably evoke ingress at latencies of tens of milliseconds (Figure 2). This behavior comes readily to naïve, head-fixed mice that have neither undergone training nor extensive acclimation, suggesting that the reliability and stereotypy of their responses are the manifestation of an innate behavioral program.

Rodents exhibit a variety of defensive behaviors in response to innately aversive and conditioned cues, including flight, freezing, crouching, defensive threat, defensive attack, and burying of potentially threatening objects (Blanchard and Blanchard, 2008; Blanchard et al., 1986; Blanchard et al., 1998). The category of response elicited by a given cue depends on the context in which it is presented (Bolles and Collier, 1976; Bouton and Bolles, 1980; Pinel and Triet, 1978; Yilmaz and Meister, 2013), the nature of the conditioned stimulus (Karpicke et al., 1977; Pinel and Triet, 1978), and the ongoing behavioral state of the animal (Fentress, 1968a, b). We observed three stimulus-induced behaviors in this assay: flinch, ingress, and tremble. We interpret flinch in response to mild air puff as a startle response (Davis, 1984); ingress as a flight-like response, whose abrupt onset and selective release by looming visual stimuli resembles stimulus-selective flight in freely-moving animals (De Franceschi et al., 2016; Yilmaz and Meister, 2013); and tremble as freezing. Indeed, while mice typically ingress in response to CS+ presentation (Figure 5), in some cases animals also exhibit a selective tremble response to the aversive cue (Figure 5—figure supplement 2), an observation we have also made in video taken of freely moving animals during aversive conditioning (data not shown). Since mice produce these distinct behaviors (ingress, tremble) in response to an unchanging external stimulus, this assay may permit the investigation of the mechanism of action selection downstream of stimulus detection.

Novel stimuli typically trigger exploration (Berlyne, 1950) but can also evoke defensive behaviors. For instance, rats have been observed to avoid for hours an unfamiliar object placed in an otherwise familiar context (Chitty and Shorten, 1946; Shorten, 1954). Placing familiar food in a novel context inhibits feeding (novelty-induced hypophagia)—a phenomenon mitigated by anxiolytics (Dulawa and Hen, 2005). These results indicate that under some circumstances novelty may be aversive. When we presented novel, neutral odor stimuli to mice in the VBA, they initially ingressed before habituating over repeated presentations. The VBA thus can measure the transition from novelty to familiarity (Cooke et al., 2015; Groves and Thompson, 1970; Horn, 1967). Moreover, since this novelty response is stimulus-specific, the VBA may also be employed as a stimulus discrimination assay that requires neither Pavlovian nor instrumental conditioning.

The VBA is compatible with standard electrophysiological and optical methods for measuring and perturbing neuronal activity. Since it measures ingress latency at a millisecond timescale it permits alignment of, and direct comparison with, neuronal dynamics. Such alignment to sharp transitions in behavioral state has proved fruitful in primate neurophysiology: for instance, alignment to the precise time of eye saccades indicating a perceptual decision permits the investigation of the neuronal events underlying a decision process (Roitman and Shadlen, 2002).

## Acknowledgments

We are grateful to Tanya Tabachnik for the design and fabrication of the frictionless rail system; Hopi E. Hoekstra and Caroline K. Hu for helpful conversations regarding burrowing behavior, as well as for providing unpublished video of mice in the wild; Ya- tang Li, Markus Meister and Samuel G. Solomon for advice on flight-inducing visual stimuli; John P. Cunningham for advice on statistical analyses; Gary W. Johnson for precision machining; J. Andrew Miri for suggesting the use of a frictionless rail; Paul R. Stegall and Damiano Zanatto for advice on mechanical engineering; Alberto Hernandez for supplying cardboard tubes; D. Caroline Blanchard, Rui M. Costa, Walter M. Fischler, Alla Y. Karpova, John W. Krakauer, and Eve Marder for comments on the manuscript; and the 2016 Janelia Junior Scientist Workshop on Neural Circuits and Behavior.

## Author contributions

This paper is the result of a close collaboration between C.E.S. and A.J.P.F., who conceived the assay and performed the experiments. All authors analyzed the data and contributed to writing the manuscript.

## Materials and methods

### Subjects and surgery

All procedures were approved by the Columbia University Institutional Animal Care and Use Committee (protocol AC-AAAI8650). 10–17-week old male C57BL/6J mice (Jackson laboratories, Bar Harbor, ME) were fitted with a titanium head plate (27.4 mm ×9.0 mm ×0.8 mm, G. Johnson, Columbia University). Animals were anesthetized with isoflurane (3% induction, 1.5-2% maintenance) and placed within a stereotaxic frame (David Kopf Instruments, Tujunga, CA) on a feedback-controlled heating pad (Fine Science Tools, Foster City, CA). Carprofen (5 mg/kg) was administered via subcutaneous injection as a preoperative analgesic and bupivacaine (2 mg/kg) was delivered underneath the scalp to numb the area of the incision. The skull was exposed, cleaned with sterile cotton swabs and covered in a thin layer of cyanoacrylate adhesive (Krazy Glue, Elmer’s Products, Atlanta, GA). After applying a coating of adhesive luting cement (C&B-Metabond, Parkell, Inc., Edgewood, NY) onto the layer of cyanoacrylate adhesive, the titanium head plate was lowered atop the skull and secured with additional application of luting cement. The headplate was centered about the body’s anterior-posterior axis and equally spaced between bregma and lambda. For mice exposed to visual stimuli head plate position was sufficiently posterior to prevent occlusion of the visual stimuli by the head plate. Mice were allowed at least one full week and typically greater than 4 weeks to recover before any testing was performed (Table 1). All animals were singly housed on a 12 hour/12 hour light/dark cycle. All experiments were performed during the animals’ dark phase.

**Table 1.**
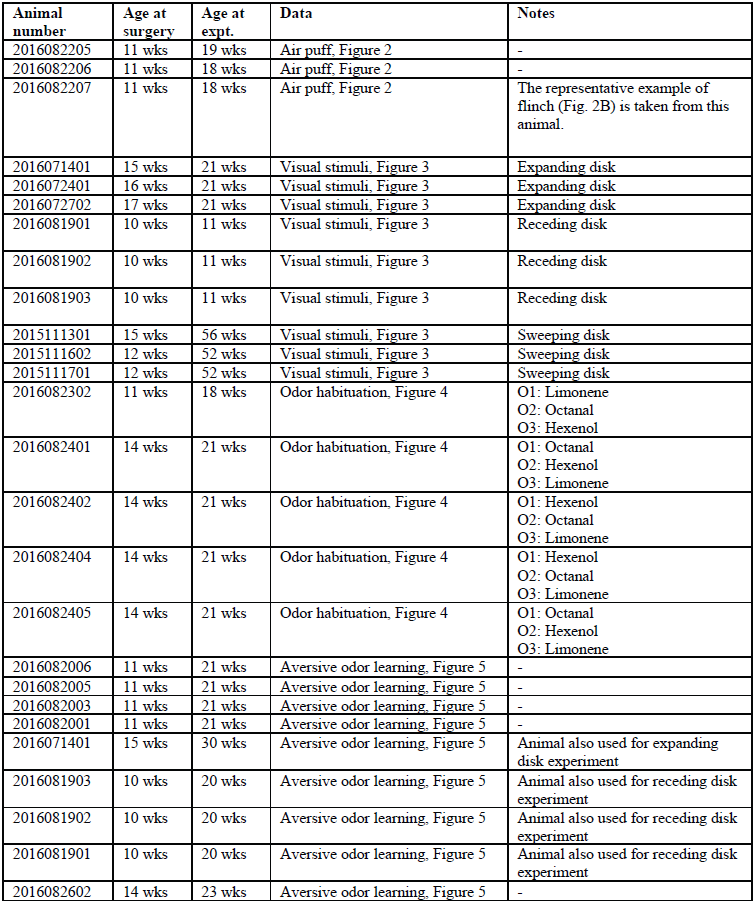
Animals used in this study

### Design of the Virtual Burrow Assay

The Virtual Burrow Assay (VBA) consists of tube enclosure (virtual burrow) constructed of a cardboard or 3D-printed polylactic acid tube (45.5-mm inner diameter, 49-mm outer diameter, 7-cm long). A 4-cm long, 0.5-mm diameter wooden rod is adhered to the front opening of the tube, 1 cm from the base, in order for animals to grip and rest their forelimbs. For the aversive learning experiments (Figure 5) the tube included a 1-cm wide extension spanning approximately 1/3 of the tube’s bottom circumference. The virtual burrow is affixed to air bearings (New Way Air Bearings, Aston, PA) that slide along two precision oriented rails, parallel to the anterior-posterior axis of the animal’s body (design and assembly: T. Tabachnik, ZMBBI Advanced Instrumentation, Columbia University; Fabrication: Ronal Tool Company, Inc., York, PA; Figure 1A, right and Figure 1B). The animal is head-fixed via custom-machined stainless steel head plate holders (G. Johnson, Columbia University) that secure the titanium headplate. The entire VBA apparatus rests atop an adjustable platform (Thorlabs, Newton, NJ) to permit precise translation of the position of the tube with respect to the head. With its head thus secured, the animal’s body rests freely within the virtual burrow, its forepaws resting on the horizontal bar placed at the burrow’s threshold, its hind limbs gripping the burrow’s interior

A linear actuator (Part number: L12-30-50-12-I, Firgelli Automations, Ferndale, WA), tethered to the virtual burrow with fishing line (0.15-mm diameter nylon tippet,4.75 pound test, Orvis, Sunderland, VT) constrains how far the animal may ingress into the burrow at any given time (Figure 1B,C). This parameter can be manually or programmatically varied over the course of the experiment. A force sensor (Futek FSH02664 load cell with Futek QSH00602 signal conditioner, Futek, Irvine, CA) reports whether, and how strongly, the animal is pulling against the linear actuator in its effort to ingress. Upon head-fixation in the VBA, mice invariably ingress as far as the linear actuator command position permits, as indicated by continuous force measured by the force sensor. When the linear actuator retracts the burrow away from the ingress position (egress position, 10-20 mm posterior to ingress position), mice resist the translation, pulling against the tether in an effort to move the burrow back up around their body. This effort typically generates between 40 and 100 g of force, corresponding in some cases to more than three times animal’s own body weight. We have not observed any mice that fail to resist retraction of the virtual burrow.

A laser displacement sensor (Part number: ILD1302-50, Micro-Epsilon, Dorfbach, Germany) is positioned so as to measure the linear displacement of the tube along its axis of motion. The laser displacement sensor is aimed at a flag affixed to the horizontal bar that joins the air bearings, whose position moves with the virtual burrow. The readout of the laser displacement sensor yields a continuous, time-dependent, one-dimensional variable. It is this quantity – how far the animal has pulled the virtual burrow around its body – that tracks ingress in response to a given cue.

For all experiments reported here the analog voltage signals from the laser displacement sensor and the force meter were acquired and digitized at 10 kHz using a Cerebus Neural Signal Processor (Blackrock Microsystems, Salt Lake City, UT).

The hardware design can be obtained in the CAD folder located at https://github.com/goatsofnaxos/VBAcmd.

### Trial structure and closed loop control

Before each trial the control system pulls the virtual burrow back to the egress position and waits until the force meter indicates that the animal has ceased to resist burrow retraction (Figure 1D). The linear actuator is then advanced to the ingress position, slackening the tether and permitting free movement of the burrow. If the animal spontaneously ingresses prior to stimulus onset, as measured by the laser displacement sensor, the trial is aborted, the burrow is again retracted to the egress position, and the sequence repeats. Once the mouse has maintained the free, egress position without attempting to ingress within a specified duration, and has maintained the standard deviation of the tube position below a user-specified threshold for a specified delay period, the stimulus is delivered. During stimulus presentation, and a set duration following stimulus offset, the control system is switched to open loop, permitting the mouse to pull the burrow up to the ingress position if it wishes.

The burrow position (measured by the laser displacement sensor), burrow force (measured by the force sensor), and the servo position (state of the linear actuator) are analog inputs to a National Instruments card with analog and digital in/out (Part number: USB-6008, National Instruments, Austin, TX). The servo position was in turn controlled by the National Instruments card (Figure 1C,D).

Prior to testing, naïve mice are head-fixed in the VBA and given 5 - 10 mn to acclimate to the contingencies in open loop (free movement of the burrow). Without exception, mice maintained the burrow in the ingress position throughout this habituation period. Then they are acclimated to the closed loop regime; after an initial period of sustained struggle to maintain the burrow in the ingress position, mice cease resisting and eventually consent to holding the burrow in the egress position even after the linear actuator has advanced, slackening the tether and granting the mouse control over the burrow. The duration of the closed-loop acclimation period varied across mice (5 - 20mn). Trial blocks begin once the animal reliably holds the burrow in the egress position for >30 sec between spontaneous ingresses. Trial initiation is delayed until after the mouse has held the burrow in the egress position with minimal movement for several seconds so as to ensure that the animal is in a comparable behavioral state prior to each trial. The control software can be obtained at https://github.com/goatsofnaxos/VBAcmd.

### Air puff stimulus

Animals were head-fixed within the VBA and permitted to acclimate to head fixation for 5 mn with the VBA on open loop. The VBA was then switched to the closed loop configuration and air puff stimuli were delivered once the animal readily gave trials. An 18-gauge, blunt syringe needle delivered air puff stimuli to generate both flinch (Figure 2D, needle tip 15 cm from nose, air pressure 2 PSI, puff duration 20 msec) and ingress (Figure 2B, needle tip 2 mm from nose, air pressure 80 PSI, puff duration 200 msec, ITI 180 sec, 15 trials per animal).

To determine latency to air puff, we subsequently measured the time between the TTL pulse controlling valve opening and the displacement of a small polystyrene weighing boat placed 2 mm distant from the blunt syringe needle (data not shown). We then subtracted the time between TTL pulse and measured displacement to determine the latency between TTL command and air puff stimulus at the nose. To account for variability in the position of the nose of the mouse with respect to the needle tip, we varied the precise location of the syringe needle over a range of distances similar to variability in distance between the syringe needle and the animal’s nose across experiments. We observed negligible variability in latencies across this distance range, demonstrating that this is not the reason why different animals exhibit different mean response latencies (Figure 2D).

For this and all experiments, a background of acoustic white noise (1000-45000 Hz; approximately 7 dB) was played throughout. The VBA apparatus was placed within a custom-made sound attenuating chamber resting on an air table (TMC, Peabody, MA). For the experiments studying responses to visual stimuli, the chamber was open to accommodate the bulk of the display screen but the lights in the room were off and the door was closed.

### Visual stimulus

For experiments examining responses to visual stimuli, animals were acclimated to head fixation within the VBA for 3 mn in the open loop configuration. Following a subsequent 10-mn habituation period with the VBA in the closed loop configuration, the animal was again permitted to freely ingress in open loop for 3-5 mn. The VBA was then returned to the closed loop configuration and once the animal did not spontaneously ingress for periods greater than 30 sec (typically after approximately 3 mn) visual stimuli were delivered.

The visual stimuli employed were based on those described in de Francheschi et al. (2016). Briefly, the stimuli were presented on a Dell 1707FP 17” LCD monitor, 1280 ×1024, 60 Hz, elevated 30 cm and centered above the animal’s head. The three stimuli, generated using the Psychophysics Toolbox 3 in MATLAB (Mathworks, Natick, MA), consisted of a black disk presented against a grey background: expanding disk (“loom”), widening from 2° to 50° over 250 msec, holding the 50° disk for 500 msec; contracting disk (“recede”), diminishing from 50° to 2° over 250 msec, holding the 2° disk for 500 msec; and sweeping disk (“sweep”), a 5° disk sweeping smoothly across the diagonal of the screen at a rate of 21°/sec. In order to permit synchronization of stimulus timing with burrow position measurement, the software controlling the visual stimulus also controlled a PWM signal (generated by an Arduino Uno, Adafruit, New York, NY; acquired as an analog voltage input digitized at 10KHz simultaneous to the position and force signals) that encoded the identity and timing of the visual stimuli.

We divided nine mice into three groups of three animals, one group per stimulus type, and presented each mouse only one of the stimulus types in a single session of five stimulus presentations separated by a 10-mn ITI. The data for each stimulus type are pooled across animals for each group.

### Odor stimulus

We used a custom built olfactometer to deliver odor stimuli. Briefly, a nose port constructed of polyether ether ketone (PEEK) was placed approximately 1 mm away from the animal’s nose. When no odor stimulus was given, the port delivered a steady stream of air (1 liter per minute, controlled by a mass flow controller, GFCS-010201 from Aalborg, Orangeburg, New York) that had bubbled through a 50-ml glass bottle containing 15 ml dipropylene glycol (DPG, Part number: D215554, Sigma-Aldrich, St. Louis, MO). To deliver an odor stimulus, a four-way valve (Part number: LSH360T041, NResearch Inc., West Caldwell, NJ) routed the air stream to exhaust, replacing it with a stream of odorized air; the odor stimulus was switched off by the four-way valve routing the odorized air back to exhaust. Monomolecular odorants (cis-3-Hexen-1-ol, catalog number W256307; (R)-(+)-Limonene, catalog number 183164; Octanal, catalog number O5608 all from Sigma-Aldrich, St. Louis, MO) were dissolved in 15 ml DPG at a concentration of 2% volume/volume in separate 50-ml glass bottles. After passing through the nose port all gas was routed to a photo-ionization detector (miniPID, Aurora Scientific, Aurora, ON, Canada) to permit constant monitoring of odorant concentration. To avoid contamination, all material in contact with the odorized air stream was constructed in either Teflon, Tefzel, or PEEK. The flow of the air and odor streams were equalized before each experiment (using mass flow meter GFMS-010786 from Aalborg, Orangeburg, New York) and the tubing carrying the two streams from the four way valve was set to equal length and impedance to minimize variation in flow rate upon switching between the air and odor streams.

### Odor habitation

For odor habituation experiments, animals (five mice) were head-fixed in the VBA and allowed to acclimate in the open loop configuration for 5 mn. The VBA was then set to closed loop for 10 - 15 mn. After habitation to the VBA, the animal was then presented with odor stimuli with the VBA in the closed loop regime. First, Odor 1 was presented 15 times. Then, a second odor, Odor 2, was introduced and the two odors were presented 15 times each, pseudo-randomly interleaved within blocks in which each of the two odors were presented in every block. Finally, a third odor, Odor 3, was added and all three odors were presented in 15 final blocks of three trials each. Each odor stimulus was presented once per block. All odor stimuli were 8 sec in duration and the ITI was 40 sec. Limonene, Octanal, and Hexenol were used as odor stimuli with different animals receiving different odors for the Odor 1, Odor 2, and Odor 3 stimuli (Table 1).

### Odor-shock conditioning and testing

*Day 1: Pre-test*. Animals used in odor-conditioning experiments were first habituated to the three odor stimuli employed (CS+: Limonene, CS-: Hexenol, O3: Octanal). Animals were placed within the VBA and acclimated to head fixation for 5 mn with the VBA in open loop, after which the VBA was switched to closed loop for 10 mn. Following the 10-mn closed loop acclimation period, the VBA was restored to the open loop configuration for 5 mn to permit the animal to freely ingress before testing, and then returned to closed loop immediately before commencing odor stimulus delivery. Odor stimuli (8-sec duration) were presented in 10 blocks of three pseudo-randomly interleaved trials (60-sec ITI) such that each stimulus was presented once per block. Following completion of the 10 stimulus blocks, animals were immediately removed from the VBA and returned to their home cage.

*Day 2: Conditioning*. Conditioning was performed one day after odor habituation. A fear conditioning box (14.2-mm wide, 16.2-mm long, 12.6-mm high, Med Associates, Fairfax, VT) was employed. Under conditions of darkness with an acoustic background of white noise, mice were placed within the fear conditioning box on the open, gloved hand of the experimenter. Once the animal had freely entered the fear conditioning box, the door was closed and the animal was allowed to acclimate for 5 mn. Eight blocks of CS+ and CS- odor stimuli were presented in pairs of pseudo-randomly interleaved trials. The odor stimuli were 10 sec in duration with a 5-mn ITI. During the final 2 sec of presentation of the CS+ stimulus only, the floor of the fear conditioning box was electrified (intensity 0.70–0.73 mAmp). Upon completion of all 8 trials, the mouse was permitted to recover for 5 mn in the fear conditioning box and then returned to its home cage.

*Day 3: Test*. One day after conditioning animals were returned to the VBA to test responses to all odor stimuli. Test was identical to pre-test except that eleven stimulus blocks were presented.

### Statistics

To determine whether responses in the VBA differed across experimental conditions, we asked whether the likelihood of ingress was larger in one condition than another. For the purposes of this test we define an ingress as a maximum change in burrow position greater than a given threshold (here 0.75 - 0.85 mm; the results of the statistical tests are robust to the choice of threshold; see Figure 5—figure supplement 1). For each condition we pooled all ingress responses across mice and used a two-proportion z-test with the null hypothesis that the probability of ingress in the tested condition was less than or equal to the probability of ingress in the other condition. One star (*) indicates p < 0.05, two stars (**) indicate p < 0.01, and three stars (***) indicate p < 0.001.

**Figure 5—figure supplement 1.**
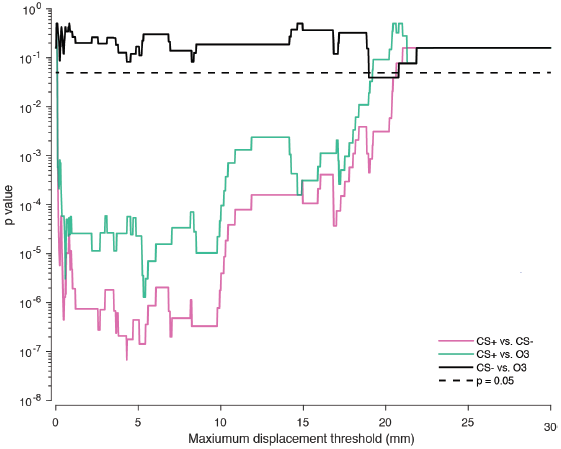
Robustness of statistical test to choice of ingress threshold. Effect of maximum displacement threshold on the two-proportion z-test p value for the pooled test data (n = 9 mice). Dashed line indicates p = 0.05.

**Figure 5—figure supplement 2.**
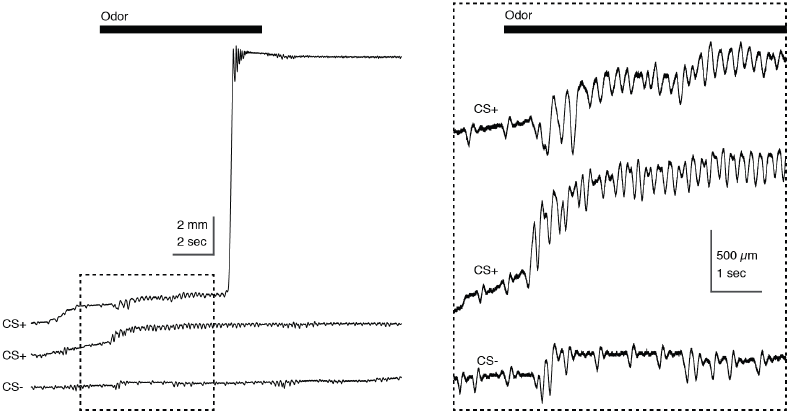
Tremble in response to aversively conditioned odor stimuli. Tube position during three individual trials recorded during Test, in response to CS+ and CS-. Odor stimulus epoch denoted by the black bar (top); dashed box at left demarcates epoch in which scaling is expanded at right. High frequency and amplitude oscillation is observed during presentation of the CS+ (top, middle) but not the CS- (bottom). Low frequency and amplitude oscillation preceding the stimulus epoch corresponds to the animal’s breathing cycle.

**Supplementary Video 1 (related to Figure 2).**
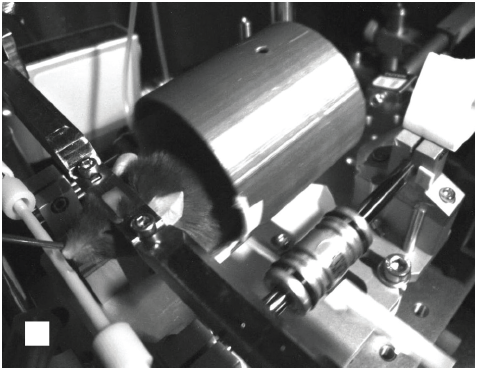
Ingress in response to strong air puff. White square: stimulus epoch (250 msec, 80 PSI, 2-mm distance from the snout).

**Supplementary Video 2 (related to Figure 3).**
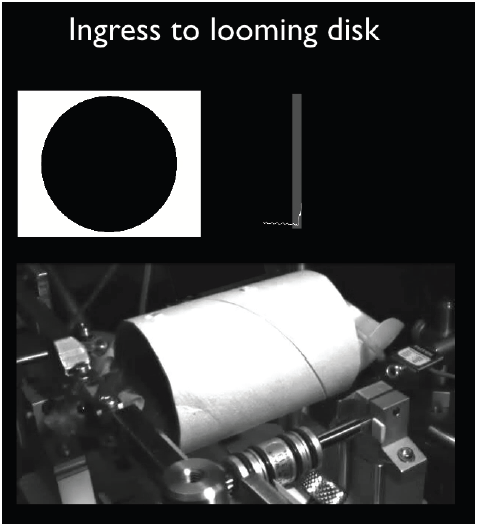
Ingress in response to visual looming stimulus. Top left panel: visual stimulus presented on a screen positioned above the animal’s head (see diagram in Figure 3A); bottom panel: video of a mouse head-fixed in the VBA; top right panel: position of the virtual burrow measured by a laser displacement sensor (grey rectangle indicates the stimulus epoch). The videos in the three panels are approximately synchronous.

